# MRI signatures of cortical microstructure in human development align with oligodendrocyte cell-type expression

**DOI:** 10.1101/2024.07.30.605934

**Authors:** Sila Genc, Gareth Ball, Maxime Chamberland, Erika P Raven, Chantal MW Tax, Isobel Ward, Joseph Yuan-Mou Yang, Marco Palombo, Derek K Jones

## Abstract

Neuroanatomical changes to the cortex during adolescence have been well documented using MRI, revealing ongoing cortical thinning and volume loss with age. However, the underlying cellular mechanisms remain elusive with conventional neuroimaging. Recent advances in MRI hardware and new biophysical models of tissue informed by diffusion MRI data hold promise for identifying the cellular changes driving these morphological observations. This study used ultra-strong gradient MRI to obtain high-resolution, in vivo estimates of cortical neurite and soma microstructure in sample of typically developing children and adolescents. Cortical neurite signal fraction, attributed to neuronal and glial processes, increased with age (mean R^2^_fneurite_=.53, p<3.3e-11, 11.91% increase over age), while apparent soma radius decreased (mean R^2^_Rsoma_=.48, p<4.4e-10, 1% decrease over age) across domain-specific networks. To complement these findings, developmental patterns of cortical gene expression in two independent post-mortem databases were analysed. This revealed increased expression of genes expressed in oligodendrocytes, and excitatory neurons, alongside a relative decrease in expression of genes expressed in astrocyte, microglia and endothelial cell-types. Age-related genes were significantly enriched in cortical oligodendrocytes, oligodendrocyte progenitors and Layer 5-6 neurons (p_FDR_<.001) and prominently expressed in adolescence and young adulthood. The spatial and temporal alignment of oligodendrocyte cell-type gene expression with neurite and soma microstructural changes suggest that ongoing cortical myelination processes contribute to adolescent cortical development. These findings highlight the role of intra-cortical myelination in cortical maturation during adolescence and into adulthood.

## 1. Context

Over the last two decades, magnetic resonance imaging (MRI) has provided invaluable insights into the developing brain, revealing ongoing cortical thinning and cortical volume loss throughout adolescence (Mills et al., 2016; Tamnes et al., 2017). However, the underlying cellular processes driving these changes are less understood. Cortical cytoarchitecture can be broadly categorised into neurites (e.g., axons, dendrites, and glial processes) and soma (e.g., neuronal, and glial cell bodies). Traditionally, synaptic pruning has been considered the primary driver of developmental changes in cortical morphology (Huttenlocher, 1979). Recent evidence, however, suggests that myelin encroachment into the grey/white matter boundary may also contribute to changes in MR contrast typically used for volumetrics, such as T_1_ (Natu et al., 2019). Developmental patterns of cortical myelination have been elucidated using magnetization transfer (MT) imaging (Paquola et al., 2019), and indirectly using T1w/T2w ratio (Grydeland et al., 2019). Despite these advances, how microstructural changes – specifically neurite and soma properties – contribute to these distinct morphological changes remains unclear.

Diffusion-weighted MRI (dMRI) is the main non-invasive MRI technique capable of probing the tissue microstructure, orders of magnitude smaller than the typical millimetre image resolution of structural MRI (Le Bihan et al., 2001). This microstructural imaging method is highly sensitive to the magnitude and direction of water diffusing within brain tissue. By employing biophysical models, it is possible to infer microscopic properties of different tissues, such as neurite signal fraction in the brain’s white matter (Alexander et al., 2019; Zhang et al., 2012). In comparison with white matter, grey matter cytoarchitecture, broadly categorized into neurites (e.g., elongated structures such as axons, dendrites and glial processes) and soma (e.g., spherical structures such as neuronal and glial cell-bodies) is more locally complex, requiring extensions of the standard models of microstructure developed for studying the white matter. Recent hardware (Fan et al., 2022; Jones et al., 2018) and biophysical modelling (Jelescu et al., 2022; Palombo et al., 2020; Tax et al., 2020) developments have enabled diffusion-weighted microstructural quantification of soma and neurite components in the cortex in vivo. The Soma and Neurite Density Imaging (SANDI; Palombo et al. (2020)), is robust, reliable (Genc et al., 2021), clinically feasible for sufficiently short diffusion times (Schiavi et al., 2023) and has been validated in ex vivo data (Ianuş et al., 2022).

Here, we examine cortical microstructural development in a sample of children and adolescents using ultra-strong gradient dMRI to identify specific changes in neurite and soma properties with age. To identify potential cellular substrates, we analyse developmental patterns of neurite and soma microstructure alongside contemporaneous trajectories of cortical cell-type specific gene expression measured in the developing cortex using data from two independent, post-mortem databases. We reveal key developmental patterns in cortical neurite and soma architecture, highlighting the contribution of active and ongoing cortical myelination processes to the macroscale changes observed in the cortex during adolescence.

## 2. Results

We apply a framework for cortical microstructure and cell-type specific gene expression analysis (Fig 1) to evaluate the cellular properties underpinning human cortical microstructural development.

**Figure 1:**
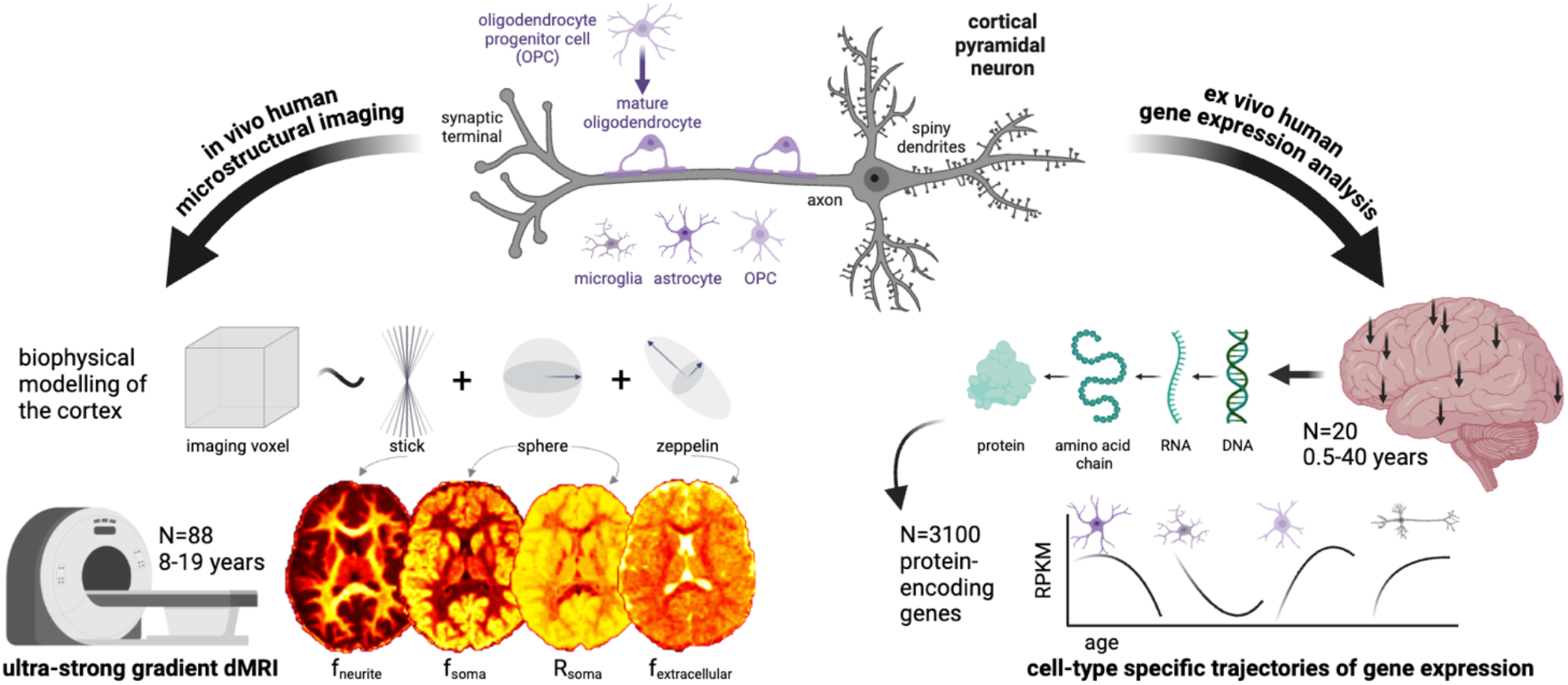
Framework for cortical microstructure and gene expression analysis. This study employs a biophysical model of cortical neurite and soma microstructure using ultra-strong gradient dMRI (Jones et al., 2018) data collected from 88 children and adolescents aged 8-19 years. Representative maps of neurite signal fraction (f_neurite_), soma signal fraction (f_soma_), apparent soma radius (R_soma_, µm) and extracellular signal fraction (f_extracellular_) are shown for one 8-year-old female participant. We also analyse two human gene expression datasets (Colantuoni et al., 2011; Li et al., 2018) to estimate cell-type specific and spatial (where arrows on brain render indicate a subset of regions sampled) gene expression profiles and examine their concordance with developmental patterns of cortical microstructure.

### 2.1. Cortical microstructure and morphology in domain-specific networks

First, we studied the repeatability of cortical microstructural estimates from the SANDI model in a sample of 6 healthy adults scanned over 5 sessions. Intra-class coefficients (ICCs) for neurite signal fraction (f_neurite_), soma signal fraction (f_soma_) and extracellular signal fraction (f_extracellular_) were very high (Fig 2c) across seven domain-specific networks (mean ICC=.97, all p<.001). Apparent soma radius (R_soma_, in µm) showed lower repeatability on average (mean ICC=.92) with lower mean repeatability driven by the limbic network (ICC=.66, p=.04).

**Figure 2:**
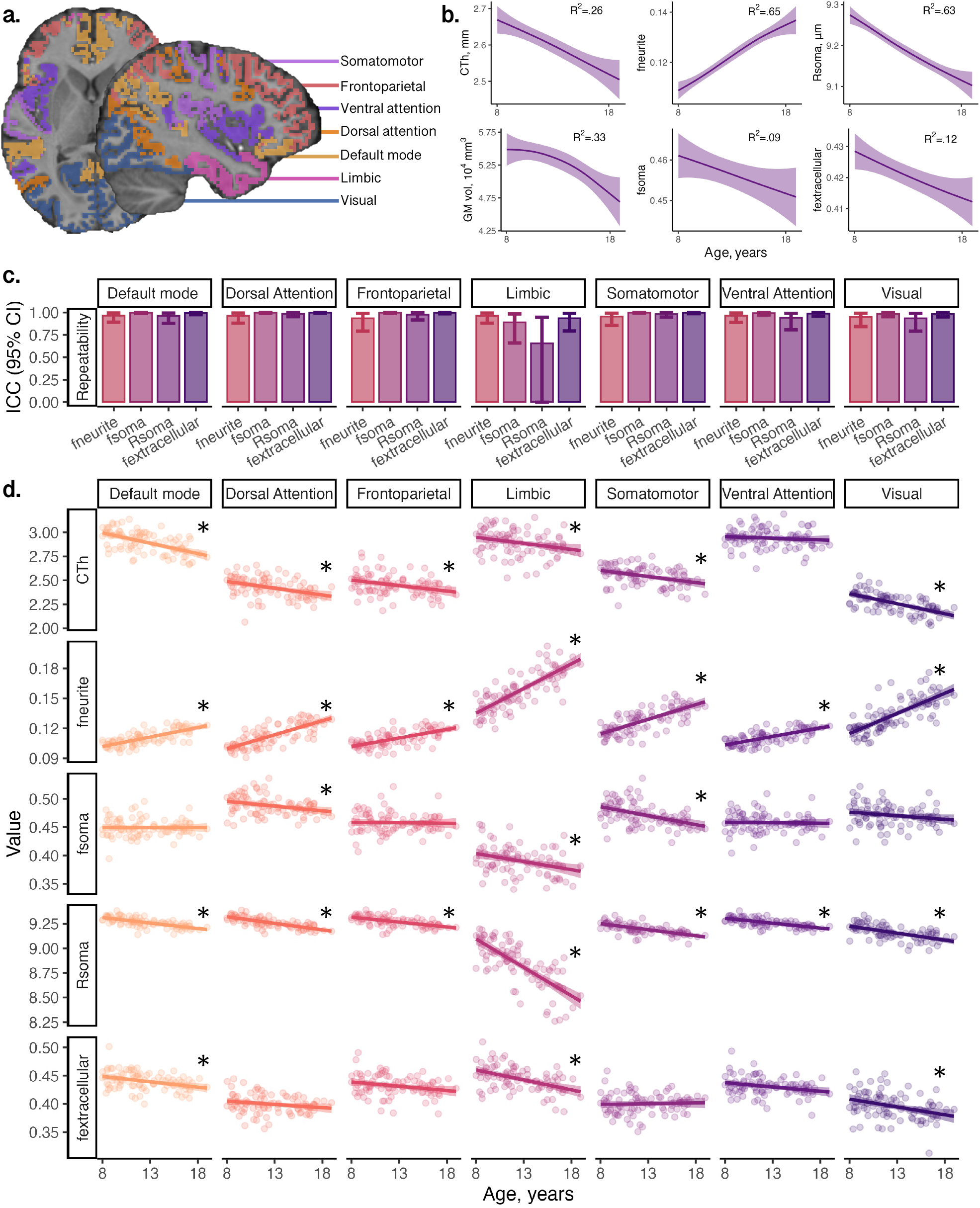
Developmental patterns of MRI-derived cortical morphology and microstructure: (a) regions in atlas used to derive domain-specific networks (Yeo et al., 2011) overlaid on a representative participant; (b) developmental patterns of cortical morphology and microstructure averaged across the cortical ribbon; (c) demonstration of high repeatability of SANDI measures in six adults scanned over 5 time-points within two weeks; (d) network-wide patterns of microstructure and morphology, indicating age-related increases in neurite fraction and reductions in cortical thickness, apparent soma radius, soma fraction and extracellular fraction. Significant age relationships (p<.005) are annotated (*). Abbreviations: CTh: cortical thickness, in mm; f_extracellular_: extracellular signal fraction; f_neurite_: neurite signal fraction; f_soma_: soma signal fraction; GM: grey matter; ICC: intra-class coefficient; R_soma_: apparent soma radius, in µm.

We then studied age-related patterns of cortical microstructure and morphology in a sample of 88 typically developing children and adolescents aged 8-19 years (Table S2). Cortical f_neurite_ and intracellular volume fraction (v_ic_; derived from the NODDI model, Zhang et al. (2012)) increased with age across all cortical networks (mean R^2^_fneurite_=.53, all networks p<3.3e-11; mean R^2^_vic_=.46, all networks p<1.6e-9) (Fig 2d, Fig S1). Orientation dispersion index (ODI; derived from the NODDI model, Zhang et al. (2012)) also increased with age across all studied networks (mean R^2^_odi_=.42, all networks p<1.9e-5). In contrast, we observed decreasing R_soma_ with age across all networks (mean R^2^_Rsoma_=.48, all networks p<4.4e-10) and f_soma_ decreased with age in the dorsal attention (R^2^_fsoma_=.12), limbic (R^2^_fsoma_=.09) and somatomotor (R^2^_fsoma_=.23), networks (all p<.002). f_extracellular_ decreased in the default mode (R^2^_fe_=.12), limbic (R^2^_fe_=.21) and visual (R^2^_fe_=.09) networks (all p<.004).

Consistent with established developmental patterns, cortical thickness and grey matter volume decreased with age (Fig 2b). The strength of these associations varied across brain networks (see Fig S1 and Table S2). Specifically, cortical thickness exhibited age-related decline in the default mode, β= -.59 [-.77, -.41], dorsal attention, β= -.40 [-.61, -.19], somatomotor, β= -.40 [-.60, -.19], and visual, β= -.61 [-.78, -.43], networks (all p<.001). Similarly, grey matter volume decreased with age in the default mode, β= -.37 [-.55, - .20], dorsal attention, β= -.34 [-.54, -.15], and visual β= -.29 [-.47, -.11], networks (all p<.002). Cortical surface area did not show significant age-related differences. The magnitude and direction of age effects across all microstructural and morphological measures are shown in Figure S1.

### 2.2. Unique sex and pubertal differences in the visual network

Sex differences in brain structure have been well reported, with pubertal onset playing a critical role in initiating developmental changes to morphology (Vijayakumar et al., 2018) and microstructure (Tamnes et al., 2018). We found that grey matter volume and surface area were higher in males than females (p<.005) across all brain networks (Figure S2), following known patterns of larger brain volume in males. We observed sex differences in only two microstructural measures, R_soma_ and fractional anisotropy (FA; derived from the diffusion tensor at b=1000s/mm^2^), in the visual network (Fig S2, S3). Females had higher R_soma_, β= -.57 [-.91, -.24], p=.001, and lower FA, β= .55, [.18, .92], p=.004, compared to males. We observed a pubertal stage by sex interaction on f_soma_, where males had lower soma signal fraction in early puberty, β= .73 [.28, 1.18], p=.002, which stabilised in late puberty. Males had lower f_extracellular_ throughout puberty β= -.74 [-1.18, -.31], p=.001.

Using an age-prediction random forest model for each microstructural measure in the visual network, we found that R_soma_ provided the most accurate age-prediction (cross-validated R^2^= .58), followed by f_neurite_ (R^2^= .56), and f_soma_ (R^2^=.28). Model fitting did not converge for f_extracellular_. NODDI measures showed R^2^_odi_=.46, and R^2^_vic_=.36. Feature importance analysis revealed that association cortices within the visual network had the highest contribution (top 5%) to age prediction (Fig 3b,c,d). Notably, region 31a (posterior cingulate cortex) consistently influenced age prediction across multiple measures, with R_soma_ contributing 63%, ODI 7% and v_ic_ 5.4%. Additional top-ranking regions included dorsal visual area, V3A (v_ic_ = 45%), lateral temporal area, TE2a (v_ic_ = 17.8 %, f_neurite_ = 5.1%), retrosplenial cortex, RSC, (vic=7.9%), auditory association area, A5 (f_neurite_=5.5%), and lateral occipital area, LO3, (f_soma_=5.4%). These regions (depicted in Fig 3c) represent cortical endpoints of developmentally sensitive tracts, identified through tractography, such as the posterior corpus callosum, cingulum, and inferior longitudinal fasciculus (Fig 3d).

**Figure 3:**
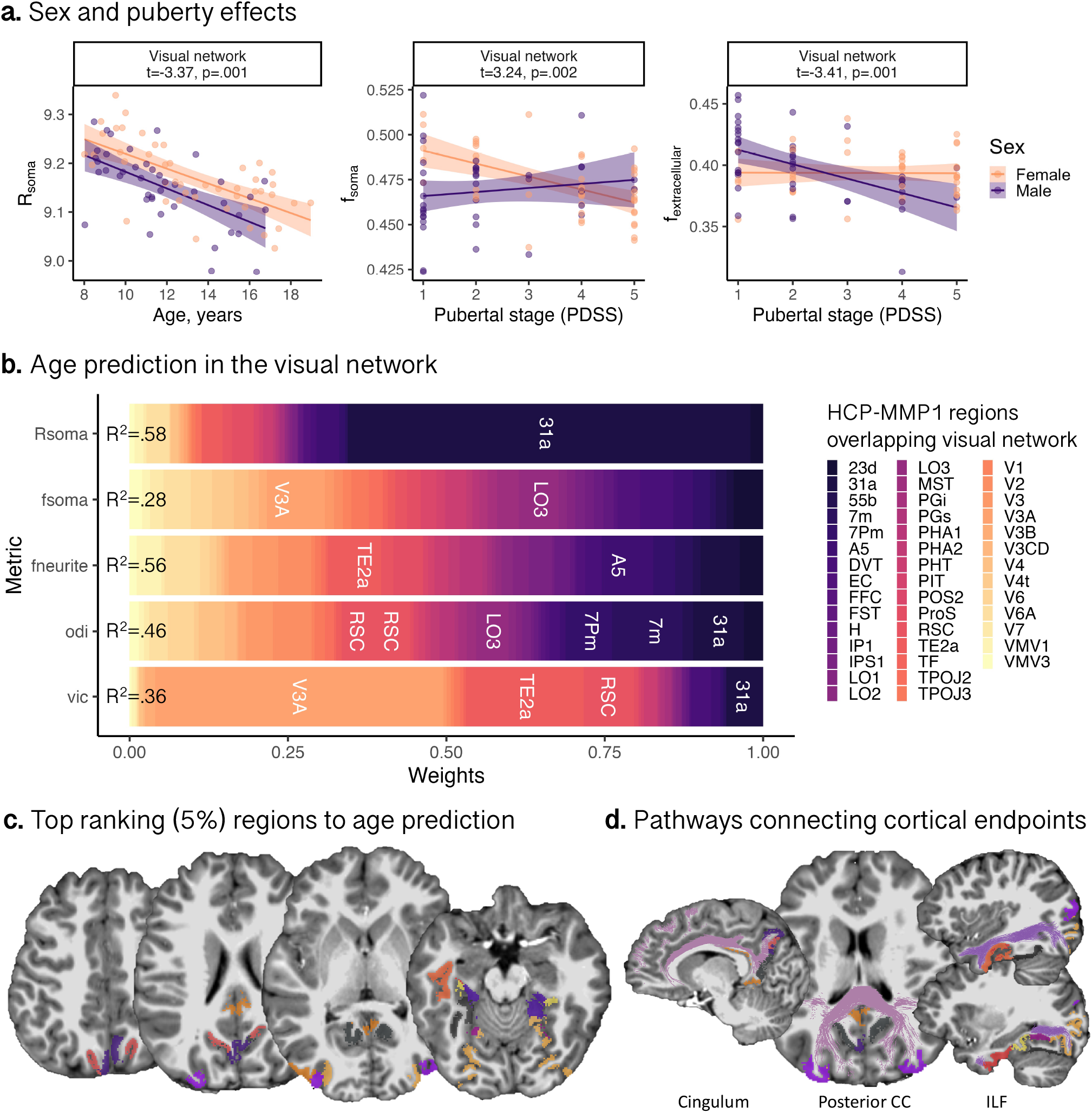
Developmental patterns of microstructure in the visual network. (a) Sex differences in apparent soma radius, and sex by puberty interactions for soma and extracellular signal fractions. (b) Feature importance of regions overlapping visual network (Glasser et al., 2016) to brain age estimation; top ranking regions with a weighting >5% (width of coloured bin) are in white text and accuracy of prediction model is represented (as R^2^) on the leftmost point of the bar plot. (c) Top ranking regions overlaid on a representative participant, coloured by labels in (b). (d) White matter pathways derived from tractography connecting cortical endpoints identified in age prediction analysis, such as the cingulum, posterior corpus callosum (CC) and inferior longitudinal fasciculus (ILF) which traverse regions in (c).

### 2.3. Contemporaneous gene expression trajectories

Using n=214 post-mortem tissue samples from the dorsolateral prefrontal cortex (DLFPC; BrainCloud; Colantuoni et al. (2011)), we identified n=2057 genes with differential expression over the lifespan (0.5 – 72 years; p_FDR_<0.05). We validated this selection in an independent RNA-seq dataset (PsychENCODE; Li et al. (2018); n=20; DLPFC samples only), identifying n=467 (22.7%) genes with significant age-associations in both datasets (age-genes; Supp Info).

We identified sets of differentially expressed genes across 7 cortical cell-types (see Methods). Mean trajectories of gene expression across the age range 0 and 30 years, averaged within each cell-type, are shown as standardized curves in Figure 4 for PsychENCODE (Fig 4a) and BrainCloud (Fig 4b) datasets. Non-normalized gene expression curves for PsychENCODE are presented in Figure S4 to aid in interpreting relative differences gene expression magnitudes. Among genes expressed in excitatory neuronal populations and oligodendrocytes, mean expression levels increased with age. In contrast, genes expressed in inhibitory neurons showed no age-related variation. Genes expressed in endothelial cells, astrocytes, microglia and OPCs, exhibited a decrease in mean gene expression with age. Overall, microglial gene expression (average log_2_RPKM =1.96) was lower compared to astrocytes (log_2_RPKM =3.70), oligodendrocytes (log_2_RPKM =3.11), OPCs (log_2_RPKM =3.01), excitatory neurons (log_2_RPKM =4.15) and inhibitory neurons (log_2_RPKM =2.94).

**Figure 4:**
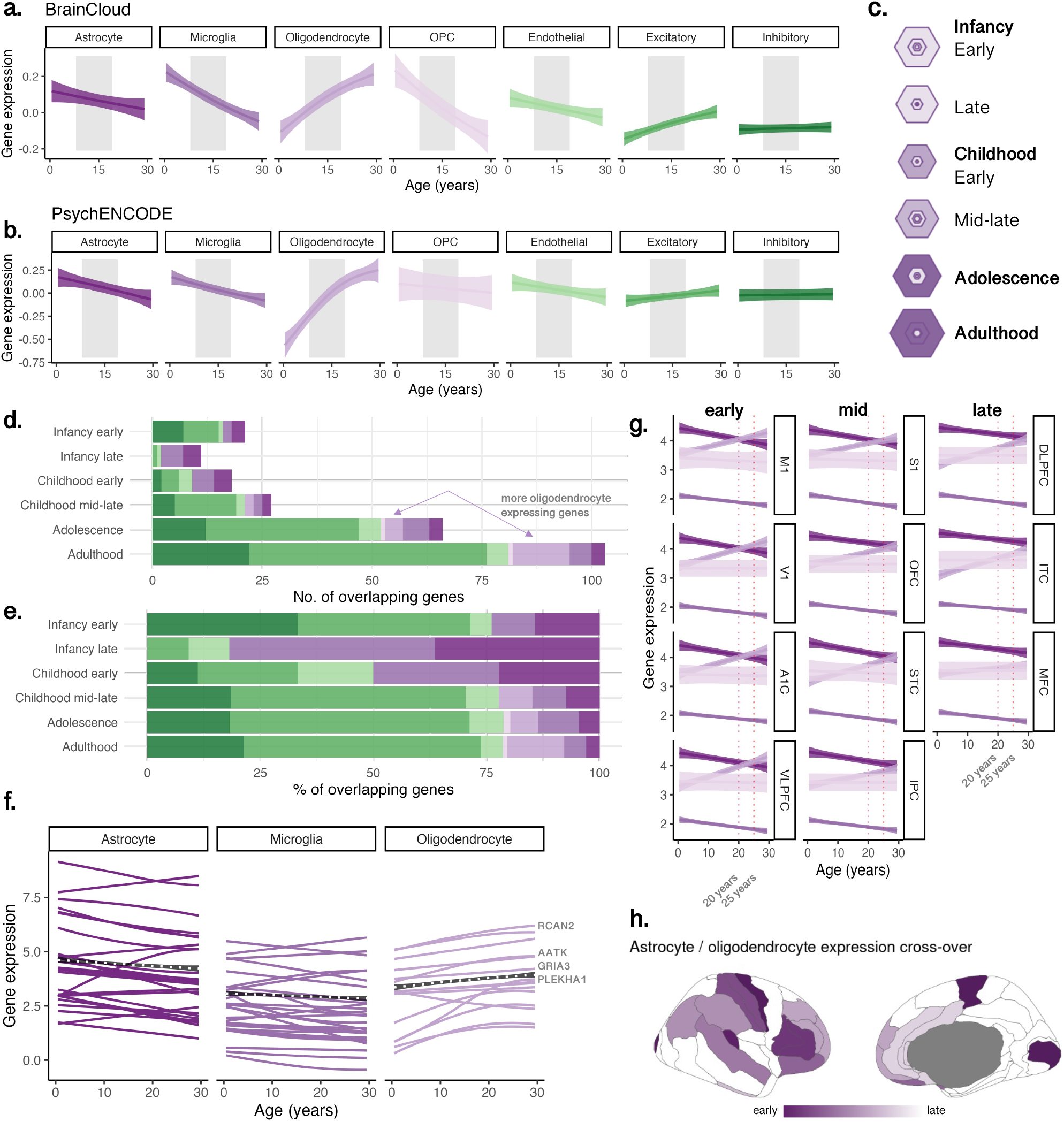
Developmental trajectories of cell-type specific gene expression. Data shown for samples aged 0-30 years from: (a) BrainCloud (Z-score), and (b) PsychENCODE (expressed in log_2_-reads-per-kilobase of transcript per million (log_2_RPKM)) datasets, demeaned to account for overall higher expression in some cell-types. Age effects were modelled in all postnatal samples to maximise sample size. Grey shaded areas highlight the age range of the microstructural imaging cohort (8-19 years) for visual comparison of developmental profiles. (c) SEA results (Xu et al., 2014) showing significant enrichment of age-related genes through adolescence and adulthood, where hexagon size scales with enrichment (overlap) of age-related genes in genes expressed by each cell type, and darker rings indicate significant associations at p_FDR_<.001 with inner rings indicating high cell specificity. Age-related genes overlapping postnatal developmental stages are shown as (d) total number of genes, and (e) proportion of genes, indicating an increase in neuronal, glial and oligodendrocyte-specific genes. (f) Trajectories of glial genes overlapping the SEA and our age-genes. (g) Regional shifts in the glial cell-type expression ratio (log_2_RPKM) across development, with the astrocyte-to-oligodendrocyte expression ratio crossing earliest at age 20 years in primary motor and visual cortices. (h) Timing of this cross-over, with darker values indicating regions with an earlier crossing point. Note that white coloured regions are not represented in the data set.

We validated the enrichment of these cell-types in these age-related genes using an independent cell-type specific expression analysis (CSEA). Significant enrichment of age-genes (n=467) was observed in cortical oligodendrocytes, oligodendrocyte progenitors, and Layer 5-6 neurons (Fig S5). These genes were prominently expressed across developmental stages in childhood adolescence, and young adulthood (Fig S6, Fig 4c). The number (Fig 4d) and proportion (Fig 4e) of age-related genes expressed by oligodendrocytes increased significantly in adolescence and young adulthood (Fig 4d,e). These included genes associated with CNS (re)myelination, RCAN2 (Huang et al., 2011), GRIA3 (Kougioumtzidou et al., 2017), and the differentiation of OPCs and oligodendrocytes, PLEHA1/TAPP1 (Chen et al., 2015); AATK/AATYK (Jiang et al., 2018).

For each cell-type, we quantified the spatiotemporal patterns of gene expression using PsychENCODE data by identifying the peak growth of expression in cell-specific genes. Oligodendrocyte gene expression peaked earliest in primary motor (M1), primary visual (V1) cortices, and latest in the medial frontal (MFC) cortex (Fig S8). A notable pattern emerged in which the peak expression of oligodendrocyte genes coincided with a shift in oligodendrocyte-to-astrocyte specific expression ratio. This shift, indicating a relative increase in oligodendrocyte over astrocyte cell-type gene expression, occurred around 20 years of age in M1 and V1, and after age 25 in DLPFC, ITC and MFC (Fig 4g,h). This sequence aligns with the known earlier myelination timing in sensorimotor cortices followed by prolonged myelination in the pre-frontal cortex into the third decade of life (Grydeland et al., 2019; Paquola et al., 2019; Sydnor et al., 2021).

### 2.4. Concordant profiles of microstructure and gene expression indicate developmental cortical myelination

To elucidate the cell-specific basis of our imaging findings, we examined neurite and soma microstructural measures in the same four frontal regions sampled in the PsychENCODE data (MFC, IFC, DLPFC, VLPFC; see Fig 5a,b) using a fine-grained parcellation of the frontal lobe. Microstructural MRI revealed regional increases in f_neurite_ and decreases in R_soma_ (Fig 5c). This pattern corresponded with increased regional oligodendrocyte cell-type gene expression profiles in the same regions over the same age period (Fig5d,e). The spatial distribution of oligodendrocyte cell-type expression was aligned with regional differences in peak growth of the neurite fraction (Fig S7). Thus, the dMRI-derived neurite signal fraction likely reflects spatiotemporal patterns of cortical myelination, matching the peak expression of oligodendrocyte-genes.

**Figure 5:**
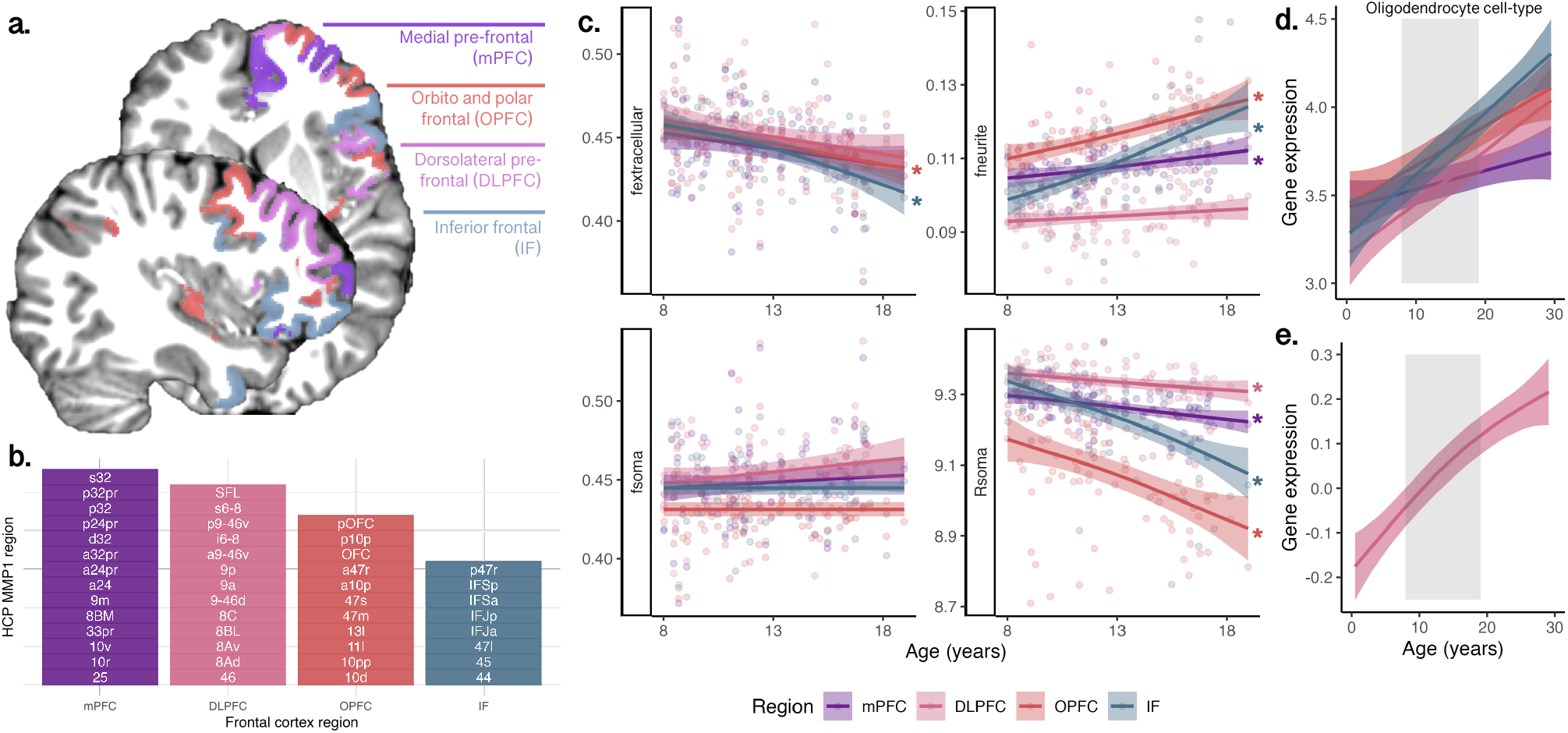
Regional variation of microstructure and gene expression in the frontal cortex. (a) Structural MRI-based segmentation of four frontal regions: medial pre-frontal cortex (mPFC); dorsolateral prefrontal cortex (DLPFC); orbito and polar frontal cortex (OPFC), and inferior frontal cortex (IF); (b) sub-regions from the HCP-MMP1 atlas (Glasser et al., 2016), which comprised the regions in (a); (c) age-related patterns of microstructural measures (*:p<.005); oligodendrocyte cell-type gene expression in (d) PsychENCODE data sampled in the same 4 frontal cortical regions as (a), and (e) BrainCloud data sampled in the DLPFC.

Numerical simulations exploring the effect of cell-type composition based on known cell counts (Keller et al., 2018) on the actual expected distribution of cell body radii within an imaging voxel reveal close correspondence between simulated and in vivo modelling results of R_soma_ (Fig S9), showing a 1% age-related decrease in both simulated and dMRI-derived data.

## 3. Discussion

We combined in vivo ultra-strong gradient dMRI with independent ex vivo gene expression analyses to map tissue microstructural architecture during human development. We now discuss each of the key findings and their implications, before summarising the strengths and limitations of our study.

### Neurite Signal Fraction Increases from Childhood to Adolescence

The neurite signal fraction, f_neurite_, attributed to elongated cortical structures (e.g., axons, processes), increased with age across the whole cortex, but peaked earliest in the visual and somatomotor networks, mirroring previous findings (Lynch et al., 2024). Intracortical myelination continues over adolescence (Bartzokis, 2012; Gibson et al., 2014; Grydeland et al., 2019; Natu et al., 2019; Whitaker et al., 2016), following a stereotyped sensorimotor-to-association (S-A) axis of development (Sydnor et al., 2023). Although dMRI is relatively insensitive to water within the myelin sheath itself, due to its short T_2_ (D. K. Jones et al., 2013), the observed increase in f_neurite_ may nevertheless reflect intra-cortical myelination. This is supported by ex vivo macaque data showing developmental increases in glial process length and complexity (Robillard et al., 2016), and an increase in the number of myelinated axons and dendrites (Fukutomi et al., 2018), which limits water exchange and leads to a greater signal contribution from inside the neurite (Jelescu et al., 2022; Olesen et al., 2022).

### Oligodendrocyte-Specific Gene Expression Increases from Childhood to Adolescence

Supporting our in vivo MRI findings, oligodendrocyte-specific gene expression increased with age (Fig 5a,b), aligning with previous observations in independent data (Paquola et al., 2019). Age-related genes were also enriched in cortical neurons (layers 5 and 6) and OPCs (Fig S6). The concordance between the human gene expression analysis (Fig S5) and the CSEA analysis based on mouse transcriptomic profiling (Fig S6; Xu et al. (2014)) indicates conservation of myelination processes via cortical oligodendrocytes. Oligodendrocyte cell turnover in the frontal cortex is dynamic, especially in adulthood, and 10 times higher in the cortex than in the white matter (Yeung et al., 2014). OPCs can generate myelinating oligodendrocytes in adulthood, even in fully myelinated regions (Richardson et al., 2011; Young et al., 2013). Importantly, oligodendrocyte function is not restricted to myelination, rather, they also perform many critical neuronal support functions beyond myelination (Bradl & Lassmann, 2010). Together our microstructural MRI and gene-expression findings converge towards increased cortical myelination through adolescence.

### Apparent Soma Radius Decreases from Childhood to Adolescence

The dMRI-derived apparent soma radius, R_soma_, decreased cortex-wide from childhood to adolescence. Neuronal soma are much larger than glial soma, measuring ∼16µm in diameter in layers 5-6 of the adult human prefrontal cortex, whereas glial soma range in diameter from 1-11µm (Rajkowska et al., 1998). Our gene expression analysis suggests specific changes in the cellular composition of the cortex with age: decreasing expression levels for astrocyte, microglia and endothelial cell-types, and (much larger) increasing expression levels for oligodendrocyte cell-types. Glial composition in the neocortex is mostly comprised of oligodendrocytes (∼75%), followed by astrocytes (∼20%) and a smaller prevalence of microglia (∼5%) (Pelvig et al., 2008). Assuming gene expression levels are proportional to cell number/density, our observations suggest a decrease in large-soma cells (e.g., endothelial), outweighed by a larger increase in small-soma cells (e.g., oligodendrocytes).

The estimated R_soma_ is dependent on the higher order moments of the soma radii distribution (i.e. skewdness and tailedness) within an MRI voxel (Olesen et al., 2022). Our own simulations of R_soma_ based on known cell composition in the human brain (Keller et al., 2018) revealed a decrease in apparent soma radii with age matching our in vivo imaging observations (i.e., a 1% decrease). This would in turn lead to a reduction in the measured dMRI signal coming from water molecules fully restricted in soma, aligning with our in vivo observations of decreasing f_soma_ with age in the limbic, somatomotor, and dorsal attention networks. It is plausible that an increase in oligodendrocyte (Peters & Sethares, 2004), not astrocyte or microglial (Robillard et al., 2016), composition could concomitantly result in a smaller average soma radii and lower soma signal fraction in the cortex through adolescence to early adulthood.

### Sex Differences in Microstructural Properties

Females have larger apparent soma radii than males, and f_soma_ and f_extracellular_ varies with pubertal stage in the visual network (Fig 3a). Pubertal hormones can stimulate apoptosis (seen in female rat visual cortex; Nunez et al. (2002)), which could explain the lower f_soma_ as puberty progresses in females. Selective neuronal cell death with unchanged glial cell number can also occur during puberty in the medial pre-frontal cortex (Markham et al., 2007; Willing & Juraska, 2015), however we did not observe any sex or pubertal differences in microstructure of the frontal cortex.

### Extracellular Signal Fraction and Myelination

In dMRI, myelin thickening can decrease the extracellular signal fraction, due to less physical space in the extracellular matrix (Jelescu et al., 2016; Derek K Jones et al., 2013). Age-related decreases in f_extracellular_ were confined to the visual network and orbito-frontal and inferior frontal cortices. Comprehensive evaluation of the myelin content is warranted to confirm the contributions of intracortical myelination to developmental changes in cortical morphology (Mancini et al., 2020).

### Spatiotemporal Patterns of Gene Expression

Peak oligodendrocyte cell-type gene expression progressed along the S-A axis, with earliest peaks in M1 and V1, and latest in MFC (Fig S8), mirroring spatial patterns of peak f_neurite_ (Fig S7). This also coincided with a relative age-related decrease in astrocyte cell-type gene expression (Fig 5g) consistent with early-life maturation of astrocytes (Bushong et al., 2004; Cahoy et al., 2008). The S-A developmental axis describes a maturation process from lower-order, primary sensory and motor (unimodal) cortices to higher-order transmodal association cortices, which support complex neurocognitive, and socioemotional functions (Margulies et al., 2016; Sydnor et al., 2021). Prolonged maturation of the pre-frontal cortex has been reported with lower myelin content in fronto-polar cortex compared with visual or somatomotor regions from childhood to adulthood (Miller et al., 2012) indicating later myelination timing. Within the frontal cortex, age-related patterns of microstructural neurite signal fraction and soma radius were prolonged in the MFC and DLPFC (Fig 5c-e). This reflects the value of estimating in vivo neurite signal fraction as these developmental hierarchies have been reproduced across various modalities (Burt et al., 2018; Gao et al., 2020; Satterthwaite et al., 2014; Sydnor et al., 2021; Vaishnavi et al., 2010; Wagstyl et al., 2015), particularly when considering the regions reaching peak maturation earliest and latest. Overall, our combined imaging genetic analyses supports the evidence of an orderly and hierarchical progression of intracortical myelination.

### Implications for Cortical Thinning

A recent study showed that cortical thinning during development is associated with genes expressed predominantly in astrocytes, microglia, excitatory and inhibitory neurons (Zhou et al., 2023). We observed faster cortical thinning of default-mode and visual networks, consistent with previous studies (Ball, Seidlitz, Beare, et al., 2020; Zhou et al., 2023). Apparent thinning may be a result of the macrostructural shift in the boundary between grey matter and white matter, in this scenario due to myelin encroachment into the cortex (Mournet et al., 2020; Natu et al., 2019). The microstructural composition of the grey matter itself may be better studied by the biophysical models used here.

### Clinical implications

Cortical morphology and myelination abnormalities are linked to various neuropsychiatric disorders (Chen et al., 2024) including schizophrenia (Alexander-Bloch et al., 2014; Wannan et al., 2019) which is characterised by deficiencies in myelination and oligodendrocyte production (Davis et al., 2003; Katsel et al., 2005). Schizophrenia patients exhibit downregulation of myelination-related genes (Tkachev et al., 2003) and post-mortem studies have shown reduced oligodendrocyte density in layer 5 of dorsolateral prefrontal cortex compared to healthy controls (Kolomeets & Uranova, 2019). Additionally, young children with autism show age-related deficits in cortical T1w/T2w ratios (Chen et al., 2022). Given these findings, quantifying cortical microstructure in such clinical cohorts is crucial, especially with adaptations towards clinically feasible acquisition protocols (Barakovic et al., 2024; Margoni et al., 2023; Schiavi et al., 2023).

### Strengths and limitations

Several methodological advancements have advanced the understanding of underlying compositional changes to cortical microstructure across development in our study. Using in vivo microstructural imaging with ultra-strong gradients (G_max_=300 mT/m; Jones et al. (2018)), we achieved sensitivity to micrometer-level imaging contrast with significant SNR improvements over clinical MRI scanners (Raven et al., 2023). Although we used a specialised system, recent advancements have enabled these measurements on more accessible, lower-gradient strength MRI systems (e.g. G_max_≥80mT/m; Schiavi et al. (2023)). Combined with two ex vivo gene expression data sets sampled from the human brain, we provide compelling evidence in favour of a framework for monitoring intra-cortical cellular composition in vivo. Further work should evaluate in vivo imaging acquisition techniques and models that account for water exchange, which can influence biophysical modelling of grey matter compartments.

Our observation of oligodendrocyte-specific gene expression increasing towards adulthood indicates the value of imaging a broader age range of young adults to fully assess trajectories of in vivo microstructural properties. It is also important to recognise that gene expression patterns do not necessarily correlate with cellular density. Histopathological confirmation is needed to verify cell size and density with biophysical signal fractions, as well as their relevancy to functional gene expression patterns.

Overall, our study provides novel *in vivo* evidence of distinct developmental differences in neurite and soma architecture, aligning with cell-type specific gene expression patterns observed in ex vivo human data. This provides a window into the role of intracortical myelination through adolescence, and how it shapes the developmental patterns of cortical microstructure in vivo.

## 4. Methods

### 4.1. Imaging set

#### 4.1.1. Participant characteristics

We included a sample of 88 typically developing children aged 8-19 years recruited as part of the Cardiff University Brain Research Imaging Centre (CUBRIC) Kids study. The study was approved by the School of Psychology ethics committee at Cardiff University. Participants and their parents/guardians were recruited via public outreach events. Written informed consent was obtained from the primary caregiver of each child participating in the study, and adolescents aged 16-19 years also provided written consent. Children were excluded from the study if they had non-removable metal implants or reported history of a major head injury or epilepsy.

We administered a survey to parents of all participants, and to children aged 11-19 years. The Strengths and Difficulties Questionnaire (SDQ) was used to assess emotional/behavioural difficulties (Goodman, 1997). The Pubertal Development Scale (Petersen et al., 1988) was used to determine pubertal stage (PDSS; Shirtcliff et al. (2009)). Additionally, we measured each child’s height and weight to calculate their Body-Mass index (BMI) (kg/m^2^).

All children and adolescents underwent in-person training to prepare them for the MRI procedure using a dedicated mock MRI scanner. This protocol was 15-30 minutes long, and designed to familiarise them to the scanner environment, to minimize head motion during the scan. All procedures were completed in accordance with the Declaration of Helsinki.

**Table 1:** Characteristics of in vivo imaging cohort.

| Measure | Summary statistics |  |  | Age relationship* |  |
| --- | --- | --- | --- | --- | --- |
|  | Mean | SD | Range | R<sup>2</sup> | p-value |
| Age, years<sup>†</sup> | 12.56 | 2.94 | 8.0 – 19.0 |  |  |
| Pubertal stage (PDSS)<sup>†</sup> | 2.89 | 1.50 | 1 – 5 | .72 | <.001 |
| SDQ, total score<sup>†</sup> | 6.45 | 3.90 | 0 – 19 | .01 | .60 |
| Body mass index, $\text{kg/m}^2$<sup>‡</sup> | 19.29 | 3.25 | 13.7 – 29.2 | .25 | <.001 |
| FSIQ<sup>^</sup> | 108 | 12.6 | 86 – 145 |  |  |

<sup>†</sup> Full sample: N=88 (42 males, 46 females)
<sup>‡</sup> Subsample: N=79 (40 males, 39 females)
<sup>^</sup> Subsample: N=48 (23 males, 25 females)
\*Age relationships determined by linear regression.

#### 4.1.2. Acquisition and processing

Discovery data: Participants aged 8-19 years (N=88, mean age=12.6 years, 46 female) underwent MRI on a 3T Siemens Connectom system with ultra-strong (300 mT/m) gradients. Structural T_1_-weighted (voxel-size=1×1×1mm^3^; TE/TR=2/2300 ms) and multi-shell dMRI (TE/TR=59/3000 ms; voxel-size=2×2×2 mm^3^; Δ = 23.3 ms, δ = 7 ms, b-values = 0 (14 vols), 500, 1200(30 dirs), 2400, 4000, 6000 (60 dirs) s/mm^2^) data were acquired.

Repeatability data: Six healthy adults aged 24-30 years (3 female) were scanned five times in the span of two weeks (Koller et al., 2021) on the same Connectom system. Multi-shell dMRI data were collected as above, with an additional 20 diffusion directions acquired at b=200 s/ mm^2^.

Pre-processing of dMRI data followed steps interfacing tools such as FSL (Smith et al., 2004), MRtrix3 (Tournier et al., 2019), and ANTS (Avants et al., 2011) as reported previously (Genc et al., 2020). Briefly, this included denoising, and correction for drift, motion, eddy, and susceptibility-induced distortions, Gibbs ringing artefact, bias field, and gradient non-uniformities. For each subject, the soma and neurite density imaging (SANDI) compartment model was fitted (Palombo et al., 2020) to dMRI data using the SANDI Matlab Toolbox v1.0, publicly available at https://github.com/palombom/SANDI-Matlab-Toolbox-v1.0, to compute whole brain maps of neurite, soma and extracellular signal fraction (f_neurite_, f_soma_, f_extracellular_ = 1 – f_neurite_ - f_soma_); the soma radius (R_soma_, in µm); and the extracellular and intra-neurite axial diffusivities (D_e_ and D_in_, respectively, in µm^2^/ms) (Fig 1). To put our results in context with previous studies, the neurite orientation dispersion and density imaging (NODDI) model (Zhang et al., 2012) was fitted to all b-values using the NODDI Matlab toolbox, publicly available at http://mig.cs.ucl.ac.uk/index.php?n=Tutorial.NODDImatlab, to estimate the intra-cellular volume fraction (v_ic_) and orientation dispersion (OD) and diffusion tensor imaging (DTI) metrics were estimated using the b=1000 s/mm^2^ shell (Fractional anisotropy (FA); mean diffusivity (MD, in s/mm^2^).

T1-weighted data were processed using FreeSurfer version 6.0 (http://surfer.nmr.mgh.harvard.edu) and post-processed to obtain network-level (N=7 ROIs; Yeo et al. (2011)) and fine-grained cortical parcellations (N=360, HCP-MMP1; (Glasser et al., 2016)). Follow-up analyses using fine-grained HCP-MMP1 parcellations in visual and frontal cortices were performed based on a priori hypotheses of earlier maturation of visual (Natu et al., 2019) and later maturation of frontal (Robillard et al., 2016) cortices, as well as for comparison with gene expression data sampled from multiple regions in the frontal cortex. Morphological measures including cortical thickness (CTh, mm), surface area (SA, mm^2^), and grey matter volume (GMvol, mm^3^) were computed at the whole brain, and parcel level. The analysis framework is detailed in Figure 1 and networks studied are depicted in Fig 2a.

### 4.2. Cortical gene expression set

Pre-processed, batch-corrected and normalised microarray and bulk RNA-seq data from postmortem human tissue samples were obtained from the BrainCloud (Colantuoni et al., 2011) (n=214; aged 6mo – 78.2y; 144 male; postmortem interval [PMI] = 29.96 [15.28]; RNA integrity [RIN] = 8.14 [0.83]) and PsychENCODE (n=20; 6mo-40y; 10 male; PMI = 17.85 [6.75]; RIN = 8.45 [0.79]) projects, respectively (Li et al., 2018). The cortical regions sampled are summarised in Table S1. Tissue was collected after obtaining parental or next of kin consent and with approval by the institutional review boards at the Yale University School of Medicine, the National Institutes of Health, and at each institution from which tissue specimens were obtained. Tissue processing is detailed elsewhere (Ball, Seidlitz, O’Muircheartaigh, et al., 2020; Li et al., 2018). Gene expression for PsychENCODE was measured as rates per kilobase of transcript per million mapped (RPKM). Gene expression for Braincloud was preprocessed and normalized following data cleaning and regressing out technical variability (see https://www.ncbi.nlm.nih.gov/geo/query/acc.cgi?acc=GSE30272).

Genes were initially filtered to include only protein-coding genes expressed in cortical cell types (n=3100, Ball, Seidlitz, O’Muircheartaigh, et al. (2020)). Using a database of single-cell RNA-seq studies, we identified genes differentially expressed across major cortical cell types (excitatory and inhibitory neurons, oligodendrocytes, oligodendrocyte precursor cells [OPCs], microglia, astrocytes, and endothelial cells (Ball, Seidlitz, Beare, et al., 2020)).

### 4.3. Statistical analyses

#### 4.3.1. In vivo imaging

We used linear regression to test for main effects of age and sex, puberty, and sex by puberty interactions. To identify the most parsimonious model and to avoid over-fitting, we used the Akaike Information Criterion (AIC) (Akaike, 1974), selecting the model with the lowest AIC. Individual general linear models were used to determine age-related differences in cortical thickness and microstructural measures in all seven Yeo networks. Evidence for an association was deemed statistically significant when p < .005 (Benjamin et al., 2018). Results from linear models are presented as the normalized coefficient of variation (β) and the corresponding 95% confidence interval [lower bound, upper bound]. We also report the adjusted correlation coefficient of the full model (R^2^).

To identify important regions that contribute to age-related differences in all the studied microstructural measures, we performed age-prediction using a random forest regressor (5-fold cross-validation) for age prediction with PyCaret (www.pycaret.org). For each microstructural measure, we randomly split the data into training and validation sets using an 80-20 ratio (total N=88: 70 training; 18 testing). Then, we performed feature scaling to ensure that all input variables (for each HCPMMP1 ROI) were on a similar scale prior to model fitting. The performance of the model was evaluated on the validation dataset. Finally, the features with the largest weight coefficients were extracted to identify specific cortical regions where variance in cortical microstructure was associated with age-related changes.

#### 4.3.2. Gene expression profiles

To identify genes differentially expressed over age (p_FDR_<.05), we modelled age-related changes in normalised expression in all available postnatal tissue samples using nonlinear generalised additive models with thin plate splines (k=5) (Wood, 2003) in R.

##### BrainCloud

The relationship between normalised gene expression and age was modelled with a nonlinear general additive model (GAM) using a penalised thin-plate spline with a maximum 5 knots:

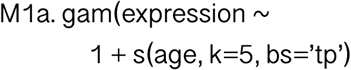

Note that the available BrainCloud data are already preprocessed to remove variance due to batch and sample effects (see https://www.ncbi.nlm.nih.gov/geo/query/acc.cgi?acc=GSE30272).

##### PsychENCODE

We repeated the above models, now with a measure of RNA integrity (RIN) as a confounder, and gene expression defined as log_2_(RPKM). First, we included region as an additional factor to account for spatial variation across the cortex and included donor ID as a random effect to account for repeated samples from the same specimen.

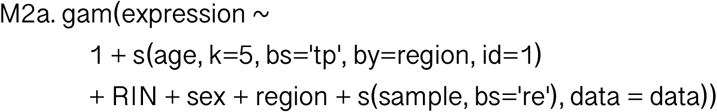

Then, we analysed data only in the DLPFC, for comparison with the BrainCloud geneset.

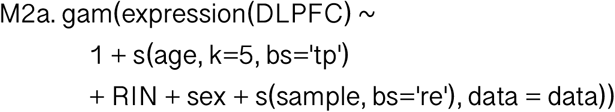

We calculated measures of goodness of fit using Akaike Information Criterion (AIC) and Bayesian Information Criterion (BIC) for all gene models.

Using a set of independent single-cell RNA studies of the human cortex (see Ball, Seidlitz et al. (2020) for details), we identified genes exhibiting differential expression across various cortical cells-types, including excitatory neurons, inhibitory neurons, oligodendrocytes, microglia, astrocytes, and endothelial cells. We then compiled gene lists for each cell-type, comprising genes that are both differentially expressed by that cell-type, and <u>uniquely</u> expressed by that cell-type. Mean trajectories across all cortical regions sampled were computed for each cell-type.

After identifying age-related genes, we entered our list to an independent cell-type specific expression analysis (CSEA; Xu et al. (2014)) to elucidate: 1) if genes were enriched for specific cell-types, and 2) in which developmental period was gene expression highest.

#### 4.3.3. Simulations

We performed numerical simulations using realistic cell counts to explain the observed trends in R_soma_ derived from in vivo dMRI data. We modelled the variability in cell body sizes within an MRI voxel by generating distributions of radii for microglia, astrocytes, oligodendrocytes, neurons, and endothelial cells. For each cell-type, we assumed the observed age-related slope of gene expression was proportional to the number of cells within an MRI voxel. Based on realistic cell counts outlined in Keller et al. (2018), we set the number of cells in mm^3^ as follows: N_micro_ = 6,500; N_astro_ = 15,700; N_oligo_ = 12,500; N_neuro_ = 92,000; N_endo_ = Nneuro*.35 (Ventura-Antunes et al., 2022). For each cell type, we generated random samples of radii based on the specified cell counts assuming a Gaussian distribution with cell-type specific baseline mean and standard deviation: microglia = 2.0±0.5 µm; astrocytes and oligodendrocytes = 5.5±1.5 µm; neurons = 8.0±2.0 µm for neurons and 9.0±0.5 µm for endothelial. The resulting radii were concatenated to form a comprehensive distribution and the MR apparent soma radius R_soma_ estimated as 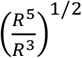 as per Olesen et al. (2022).

## Abbreviations

A1C: primary auditory cortex
AIC: Akaike information criterion
CSEA: cell-specific expression analysis
DLPFC: discovery rate
f_extracellular_: extracellular signal fraction
f_neurite_: neurite signal fraction
f_soma_: soma signal fraction
v_ic_: intracellular volume fraction
IPC: inferior parietal cortex
ITC: inferior temporal cortex
M1: primary motor cortex
MD: mean diffusivity
MFC: medial frontal cortex
MRI: magnetic resonance imaging
mRNA-seq: mRNA sequencing
NODDI: neurite orientation dispersion and density imaging
ODI: orientation dispersion index
OFC: orbitofrontal cortex
OPC: oligodendrocyte precursor cell
RIN: RNA integrity number
RNA-seq: RNA sequencing
ROI: regions of interest
RPKM: reads per kilobase of transcript per million mapped reads
S1: primary sensory cortex
SANDI: Soma and Neurite Density Imaging
STC: superior temporal cortex
V1: primary visual cortex
VLPFC: ventrolateral prefrontal cortex

## 5. Acknowledgements

The authors would like to thank the families that participated in this study for their generous contributions. We would also like to thank Umesh Rudrapatna and John Evans for their assistance with acquisition protocols and Greg Parker for assistance with the image pre-processing pipeline. Figure 1 was created with BioRender.com with elements from Macrovector via freepik.

